# Inhibition of NLRP1 Inflammasome Activation by Tyrosine Kinase Inhibitors Restores Erythropoiesis in Diamond-Blackfan Anemia Syndrome

**DOI:** 10.1101/2025.02.20.639294

**Authors:** Juan M. Lozano Gil, Lola Rodríguez-Ruiz, Manuel Palacios, Jorge Peral, Susana Navarro, José L. Fuster, Cristina Beléndez, Andrés Jérez, Laura Murillo-Sanjuán, Cristina Díaz-de-Heredia, Guzmán López-de-Hontanar, Josune Zubicaray, Julián Sevilla, Francisca Ferrer-Marín, María P. Sepulcre, María L. Cayuela, Diana García-Moreno, Alicia Martínez-López, Sylwia D. Tyrkalska, Victoriano Mulero

**Affiliations:** Departamento de Biología Celular e Histología, Facultad de Biología, Universidad de Murcia, 30100 Murcia, Spain; Instituto Murciano de Investigación Biosanitaria (IMIB-Pascual Parrilla), 30120 Murcia, Spain; Centro de Investigación Biomédica en Red (CIBERER), ISCIII, 28029 Madrid, Spain; Centro de Investigaciones Energéticas, Medioambientales y Tecnológicas (CIEMAT), 28040 Madrid, Spain; Unidad de Terapias Avanzadas, IIS-Fundación Jiménez Díaz (IIS-FJD, UAM), Madrid, Spain; Sección de Oncohematología Pediátrica, Servicio de Hematología, Hospital Clínico Universitario Virgen de la Arrixaca, 30120 Murcia, Spain; Hospital General Universitario Gregorio Marañón, Instituto Investigación Sanitaria Gregorio Marañón, Facultad de Medicina Universidad Complutense, ERN-EuroBloodNet, 28007 Madrid, Spain; Departamento de Hematología y Oncología Clínica, Hospital General Universitario Morales Meseguer, 30008 Murcia, Spain; División de Hematología y Oncología Pediátrica, Hospital Universitario Vall d’Hebron, 08035 Barcelona, Spain; Pediatric Hematology and Oncology Department, Hospital Infantil Universitario Niño Jesús, 28009 Madrid, Spain; Universidad Católica San Antonio (UCAM), 30107 Murcia, Spain

**Keywords:** NLRP1, inflammasome, tyrosine kinase inhibitors, hematopoietic stem cells, zebrafish, drug repurposing, Diamond-Blackfan anemia syndrome

## Abstract

Diamond-Blackfan Anemia Syndrome (DBAS) is characterized by impaired erythropoiesis due to dysfunctional ribosome biogenesis and aberrant cellular signaling. Here, we investigate how ribosomal stress-induced activation of the NLRP1 inflammasome modulates erythroid differentiation in DBAS. We demonstrate that FDA/EMA-approved tyrosine kinase inhibitors (TKIs) effectively mitigate defective erythropoiesis in Diamond-Blackfan anemia syndrome (DBAS) by inhibiting NLRP1 inflammasome activation. Specifically, nilotinib enhances erythroid differentiation in K562 cells through suppression of the ZAKα/P38/NLRP1/CASP1 axis, leading to increased GATA1 protein levels and upregulation of key erythroid genes involved in iron acquisition, hemoglobin synthesis, and erythrocyte structure. These effects were validated in human CD34^+^ hematopoietic stem and progenitor cells (HSPCs) and zebrafish models, where nilotinib, along with other TKIs (imatinib, dasatinib, and bosutinib), promoted erythropoiesis at the expense of myelopoiesis and reduced caspase-1 activity. Importantly, in RPS19-deficient zebrafish and human models and HSPCs from patients with DBAS, nilotinib, imatinib and dasatinib rescued defective erythroid differentiation and restored hemoglobin levels. These findings highlight the potential of TKIs to address the erythroid defects observed in ribosomopathies like DBAS. Given the limited treatment options available for DBAS and other congenital anemias, our study provides compelling evidence for repurposing TKIs as a novel therapeutic strategy to alleviate pathological NLRP1 activation and improve erythropoiesis. This work opens new avenues for managing ribosome-related disorders and advancing personalized medicine approaches for hematopoietic diseases.

## Introduction

Initially identified as key regulators of innate immunity and inflammation, inflammasomes have recently been implicated in a broader range of biological processes, including hematopoiesis(Rodriguez-Ruiz et al., 2020). Furthermore, while the expression of inflammasome components was first reported in macrophages, granulocytes, and dendritic cells(Evavold and Kagan, 2019; Martinon et al., 2002), and subsequently in lymphocytes(Martin et al., 2016; Phan et al., 2007), emerging evidence demonstrates that hematopoietic stem and progenitor cells (HSPCs) also express inflammasome components(Lenkiewicz et al., 2019; Ratajczak et al., 2020; Rodriguez-Ruiz et al., 2023; Tyrkalska et al., 2019). HSPCs not only express inflammasome components but also rely on these components, including the well-characterized NLR family pyrin domain containing 3 (NLRP3) inflammasome, for the regulation and modulation of hematopoiesis. The effects of NLRP3 on HSPCs are mediated through both direct and indirect pathways. On one hand, NLRP3 activation influences the development and proliferation of HSPCs within the hematopoietic tissue(Frame et al., 2018; Ratajczak et al., 2020). On the other hand, it regulates their migration from the bone marrow into the peripheral blood in response to pharmacological mobilization or stress(Lenkiewicz et al., 2019; Ratajczak et al., 2019). Additionally, NLRP3 plays a role in guiding the reintegration of HSPCs into the bone marrow following transplantation and contributes to the aging process of these cells(Adamiak et al., 2020). Moreover, inflammasome activation within the bone marrow microenvironment is thought to modulate the balance of distinct hematopoietic cell populations. Although there is no direct evidence that NLRP3 in HSPCs regulates their differentiation, the involvement of downstream inflammasome components, such as apoptosis-associated speck-like protein containing a caspase recruitment domain (ASC) and caspase-1 (CASP1), in this process suggests that other NLRs may be implicated instead(Tyrkalska et al., 2019). Regardless, dysregulation of the NLRP3 inflammasome is closely associated with impaired myelopoiesis and lymphopoiesis, contributing to the development of hematologic disorders, including myelodysplastic syndromes, myeloproliferative neoplasms, leukemia, and graft-versus-host disease after transplantation(He et al., 2016).

Our group was the first to report that the canonical inflammasome of HSPCs orchestrates the erythroid vs. myeloid lineage decision by cleaving and inactivating the master erythroid transcription factor GATA1(Tyrkalska et al., 2019). Although this mechanism is conserved in zebrafish and humans, the specific NLR sensor involved and its activation mechanism remained unknown until recently, when we demonstrated that NLRP1 orchestrates this regulation and is activated in HSPCs via phosphorylation of S107 in its linker domain by the ZAKα/P38 axis following erythroid differentiation-induced ribotoxic stress(Rodríguez-Ruiz et al., 2023). This mechanism of activation highlighted the potential of inhibiting ZAKα using FDA/EMA-approved tyrosine kinase inhibitors (TKIs), such as nilotinib, as a promising therapeutic strategy to prevent excessive or pathological NLRP1 activation in ribosomopathies, such as Diamond-Blackfan anemia Syndrome (DBAS). The primary cause of DBAS are germline heterozygous loss-of-function mutations in small- or large-subunit ribosomal protein genes, resulting in defective ribosome biogenesis and/or function. Among these, mutations in RPS19 are the most prevalent, accounting for approximately 25% of cases(Da Costa et al., 2020). In this study, we tested several TKIs for their ability to modulate NLRP1 activation and demonstrated their therapeutic efficacy in restoring erythropoiesis in models of DBAS, paving the way for their repurposing in the treatment of ribosomopathies.

## Materials and Methods

### Zebrafish

Zebrafish (*Danio rerio* H.) were obtained from the Zebrafish International Resource Center and mated, staged, raised and processed as described(Westerfield, 2000). The lines *Tg(lyz:DsRED2)^nz50^* (Hall et al., 2007), *Tg(gata1a:DsRed)^sd2^* (Traver et al., 2003) and casper (*mitfa^w2/w2^; mpv17^a9/a9^*)(White et al., 2008) were previously described. The *spint1a^hi2217Tg/hi2217Tg^* line was isolated from insertional mutagenesis screens (Amsterdam, Burgess et al., 1999). The experiments performed comply with the Guidelines of the European Union Council (Directive 2010/63/EU) and the Spanish RD 53/2013. The experiments and procedures performed were approved by the Bioethical Committees of the University of Murcia (approval number #669/2020).

### CRISPR-Cas9 injection in zebrafish

Negative control crRNA (catalog no. 1072544, crRNA standard) and crRNA for *rps19*, *nlrp1*and *zaka* (**Table S1**), and tracrRNA were resuspended, duplexed and injected as described previously(Rodriguez-Ruiz et al., 2023). The same amounts of gRNA were used in all experimental groups. The efficiency of each crRNA was tested by amplifying the target sequence with a specific pair of primers (**Supplementary Table 1**) and the amplicon was then analyzed by the TIDE webtool (https://tide.nki.nl/). The edition efficiency obtained was about 35% for *rps19*, and 60% for *nlrp1* and *zaka*.

### Chemical treatments of zebrafish larvae

One dpf larvae were manually dechorionated at 24 hpf and treated for 24 h by bath immersion with the TKIs nilotinib (#HY-10159, 1 µM), imatinib (#HY-15463, 1 µM and 10 µM), dasatinib (#HY-10181, 0.1 µM and 1 µM), bosutinib (#HY-10158, 0.1 µM and 1 µM) and ponatinib (#HY-12047, 0.1 µM and 1 µM), all from MedChemExpress, diluted in egg water supplemented with 0.1% DMSO. *Imaging of zebrafish larvae*

Larvae were anaesthetized in embryo medium with 0.16 mg/ml buffered tricaine and whole-body images were taken with a Leica MZ16F fluorescence stereomicroscope. The number of neutrophils (lyz^+^), erythrocytes (gata1a^+^ in the yolk sac extension) and HSPCs (runx1^+^ in the CHT) was determined by counting them visually in blinded samples(Tyrkalska et al., 2019).

### Hemoglobin staining

Three dpf zebrafish larvae were anesthetized in buffered tricaine and incubated with 0.62 mg/ml o-dianisidine at room temperature for 15-45 minutes in the dark. The reaction was monitored under a dissection microscope. Once the staining was completed, they were washed 3 times for 5 minutes each time with water. The larvae were then fixed for 2 h in 4% methanol-free formaldehyde, were rinsed with PBS containing 0.1% Tween 20 (PBST) three times and stored at 4°C until imaging with a Leica MZ16F fluorescence stereomicroscope.

### Caspase-1 Activity Assays

Caspase-1 activity was determined in larval/cell extracts with the fluorometric substrate Z-YVAD 7-Amido-4-trifluoromethylcoumarin (Z-YVAD-AFC, caspase-1 substrate VI, Calbiochem), as previously described(Angosto et al., 2012; Lopez-Castejon et al., 2008; Tyrkalska et al., 2016). Caspase-1 activity in live cells was determined using FAM FLICA Caspase-1 kit (#ICT097, Biorad) following the manufacturer’s instructions. Briefly, cells were washed twice with PBS, incubated for 45 min at 37°C with 1X FAM FLICA in PBS protected from light, washed twice with 1X Apoptosis Wash Buffer, and immediately analyzed by flow cytometry.

### CD34^+^ cells purification from human cord blood and CRISPR/Cas9 gene editing

CD34^+^ HSPCs were either enriched by immunomagnetic bead selection from donated human cord blood using an AutoMACS instrument (Miltenyi Biotec) in accordance with the manufacturer’s instructions or purchased from ZenBio (#SER-CD34-F) or StemCell Technologies (#70008). Isolated cells were culture at 37°C in 5% CO_2_ in HSPC expansion media (StemSpan SFEM II) supplemented with StemSpan CD34^+^ Expansion Supplement (#02691) (both from StemCell Technologies), with the cell concentration being maintained between 0.2 × 10^6^ and 1.0 × 10^6^ cells/ml. Human cord blood samples were provided by Centro de Transfusión de la Comunidad de Madrid under Review Board agreement (SCU IIS FJD-CTCM 2022).

The sgRNAs used for editing *RPS19* and the control gene *AAVS1* have been already described(Bhoopalan et al., 2023). RNP complex was prepared by incubating Cas9-3×NLS protein (alt-rtm s.p. hifi cas9 nuclease v3, IDT) and sgRNA (with 2′-O-methyl 3′ phosphorothioate modifications in the first and last 3 nucleotides, Synthego) at a 1:3 molar ratio for 15 minutes at room temperature. Next, 20,000 HSPCs were washed with PBS and resuspended in P3 buffer (Lonza, catalog V4LP-3002) before RNP was added for a final Cas9 concentration of 0.3 mg/mL and a total volume of 20 μL. Electroporation was performed with a Lonza 4D-nucleofector (#AAF-1003X), using program DZ100, in accordance with the manufacturer’s instructions. After RNP treatment, cells were resuspended in HSPC expansion medium and left to recover for 2 days before performing the erythroid differentiation assay (see below).

### Erythroid Differentiation Assays

K562 cells (CRL-3343; American Type Culture Collection) were maintained and subcultured as described previously(Rodriguez-Ruiz et al., 2023). Cells were pre-treated for 24 h with 0.1% DMSO alone or containing the different TKIs and then differentiated for 24 h with 50 µM hemin (#16009-13-5, Sigma-Aldrich)(Smith et al., 2000). Cells were collected at 0 and 24 h post-hemin addition, washed twice, and stored at -80 °C for further analysis.

Human cord blood CD34^+^ HSPCs were differentiated in erythroid differentiation medium [IMDM with stabilized glutamine (ThermoFisher Scientific, #12440-061), 2 % human AB plasma (SeraCare, #501973), 3 % human AB serum (Atlanta Biologicals, #S40110), 1 % Pen/Strep (ThermoFisher Scientific), 3 UI/ml heparin (Sigma-Aldrich, #H3149), 10 µg/ml insulin (Sigma-Aldrich, #I9278-5ML), 200 µg/ml holo-transferrin (Sigma-Aldrich, #T0665), 1 IU EPO (Peprotech, # 100-64), 10 ng/ml SCF (PeproTech, #300-07), 1 ng/ml IL-3 (PeproTech, #200-03)] in the presence of DMSO or 0.1 µM nilotinib for 7 days. Human cord blood CD34^+^ HSPCs at different days of differentiation were stained with antibodies that recognize cell surface markers CD235A (#349112, BioLegend) and CD71 (#334108, BioLegend) to assess erythroid differentiation by flow cytometry. All flow cytometry analyses were performed with a FACS Calibur (BD Biosciences) and analyzed with FlowJo software (Tree Star).

### Erythroid/myeloid colony formation assay by mononuclear cells from DBAS patients

Bone marrow aspirates were collected at the Hospital General Universitario José María Morales Meseguer under CEIC approval number EST: 12/16, while peripheral blood was collected at the Hospital Clínico Universitario Virgen de la Arrixaca under CEIm approval number 2020-6-6-HCUVA. The age and mutations of the patients are summarized in **Supplementary Table 2**.

BMMCs and PBMCs from DBAS patients were counted and seeded in 6-well plates at 100,000 cells/ml in methylcellulose (MethoCult™ H4431, StemCell Technologies). Cultures were maintained at 37°C, 5% CO_2_ and 95% relative humidity. Erythroid (Burst-Forming Unit-Erythroid, BFU-E) and myeloid (Colony-Forming Unit-Granulocytes/Macrophages, CFU-GM) colonies were determined 14 days after cell seeding using morphological criteria.

### Immunoblotting

K562 cells were lysed in 50 mM Tris-HCl (pH 7.5), 150 mM NaCl, 1% (w/v) NP-40 and fresh protease inhibitor (1/20, #P8340, Sigma-Aldrich). Protein quantification was performed with the BCA kit using BSA as standard(Rodriguez-Ruiz et al., 2023). The primary antibodies used were human GATA1 (#3535, Cell Signaling), human NLRP1 (#AF6788, R&D Systems), human ZAKα (#A301-993A, Bethyl Laboratories), human phosphoP38 (#MA5-15177, ThermoFisher Scientific) and ACTB-HRP (#sc-47778, Santa Cruz Biotechnology). The secondary antibodies used were anti-sheep IgG (#31480, Thermofisher) andanti-rabbit IgG (#A6154, Sigma-Aldrich).

### RNA sequencing, bioinformatic analysis and validation by RT-qPCR

Three biological replicates (15 μl at 200 ng/μl) were sent to Novogene for RNA sequencing. Sequencing was performed on an Illumina NovaSeq platform, generating 150 bp paired-end reads. Quality control and differential expression analysis were conducted by the bioinformatics service of the Novogene. Using the data provided, various representations of the differential expression analysis were generated using GraphPad. RT-qPCR was performed as described previously(Rodriguez-Ruiz et al., 2023) and normalized to the *ACTB* content in each sample using the Pfaffl method(Pfaffl, 2001). The primers used are shown in **Supplementary Table 3**.

### Statistical Analysis

Data were shown as mean ± SEM and analyzed by analysis of variance (ANOVA) and a Tukey or Bonferroni multiple range test to determine differences among groups. Differences between two samples were analyzed by Student’s *t*-test. At least three independent experiments were performed with zebrafish larvae and biochemical studies. All larvae from the different independent experiments were pooled for plotting and statistical analysis. The total larvae analyzed in each experiment is indicated in all Figures. Three independent caspase-1 activity assays were performed in all experiments using a pool of 30 larvae and one representative experiment is shown with several technical replicates. Colony and erythroid differentiation assays with primary cells were performed with two technical replicates per donor. A log rank test with the Bonferroni correction for multiple comparisons was used to calculate the statistical differences in the survival of the different experimental groups.

## Results

### Nilotinib promotes erythroid differentiation of K562 cells and CD34^+^ HSPCs

We first aimed to further understand the effect of the inhibition of the ZAKα/P38/NLRP1 axis with nilotinib on the progression of hemin-induced erythroid differentiation and GATA1 accumulation in K562 cells(Rodriguez-Ruiz et al., 2023) (**Figure 1A**). We performed a RNA-seq analysis of K562 cells upon their differentiation with hemin in the presence of nilotinib and found that it was able to induce the transcript levels of several genes encoding major factors involved in erythroid maturation and function, such as heme biosynthesis (*ALAS2*, *ALAD, FECH* and *HMBS)*, the receptor of erythropoietin (*EPOR*), structural molecules located in the plasma membrane of erythrocytes (*GYPC* and *SLC4A1*), and globin subunits (*HBA1* and *HBA2*) (**Figures 1A, 1B, Supplementary Data Set1 and Set2)**. These results were confirmed by RT-qPCR (**Figures 1C-1E)**. Curiously, nilotinib acted as a better inducer of erythroid differentiation in K562 than hemin, promoting a strong expression of the above-mentioned genes without hemin treatment (**Figures 1C-1E**).

**Figure 1.**
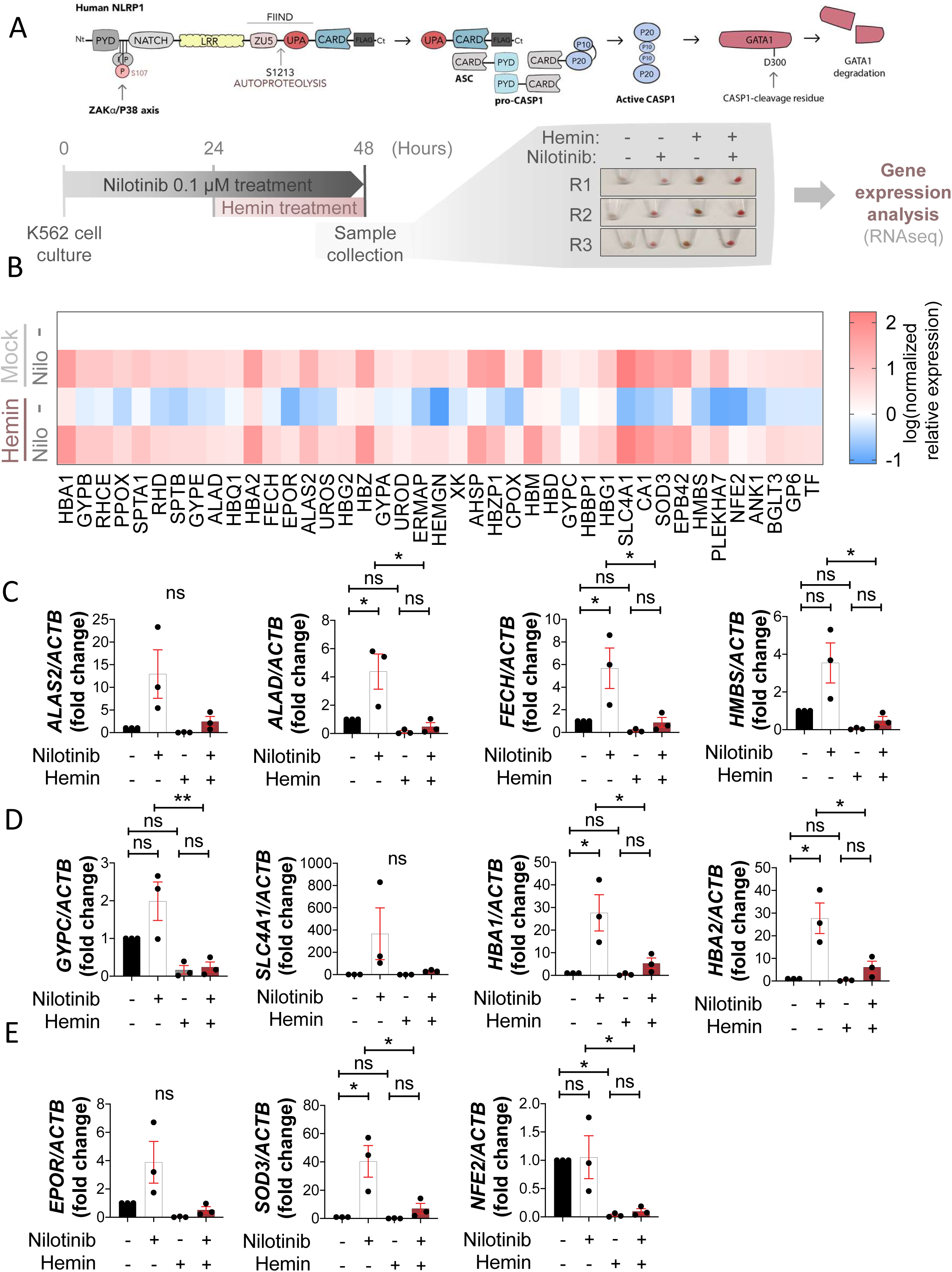
Nilotinib induces the expression of key erythroid genes in K562 cells. (A) K562 cells were pretreated with 0.1 μM nilotinib, followed by differentiation with 50 μM hemin for 24 h before transcriptomic analysis (N=3). (B) Heatmap showing the top upregulated erythroid genes upon nilotinib treatment. (C-E) RT–qPCR analysis of select upregulated erythroid genes involved in structural components of erythrocytes (C), heme biosynthesis (D), and other erythropoietic functions (E) following nilotinib treatment. Data are shown as the mean ± SEM. ns, non-significant, *p<0.05 and **p<0.01 according to a Student’s *t*-test.

The results obtained with nilotinib in K562 cells prompted us to investigate if nilotinib was also able to promote erythroid differentiation of HSPCs. With this aim, human cord blood CD34^+^ cells were cultured in differentiation media containing EPO in the presence of either DMSO or nilotinib from 3 days post-differentiation (dpd). During this process CD34^+^ cells are expected to abandon their multipotency and start expressing the receptor of transferrin (CD71) at the CFU-E stage. Then a co-expression of glycophorin A (CD235A) together with CD71 marker was expected in erythroblasts until the CD71 expression was downregulated in reticulocytes/erythrocytes. Nilotinib was able to inhibit CASP1 activity and robustly increase the transcript levels of GATA-1dependent genes and the percentage of erythroblasts (CD71^+^/CD235A^+^ cells) **(Figures 2A-2D)**, suggesting that nilotinib promoted erythropoiesis We repeated the experiments adding nilotinib at 7 dpd and, in this case, nilotinib also inhibited CASP1 activity but the effect in promoting differentiation was weaker than in the previous experiment **(Figures 2E-2H)**. Importantly, the inhibition of CASP1 activity by nilotinib in both experiments confirmed that it was inhibiting the ZAKα/P38/NLRP1 inflammasome. All these results suggest that nilotinib induces the progression of erythroid differentiation, which may be of clinical interest since some types of congenital anemia do not only show decreased number of HSPCs but also defective erythropoiesis, for example DBAS (Da Costa et al., 2020).

**Figure 2.**
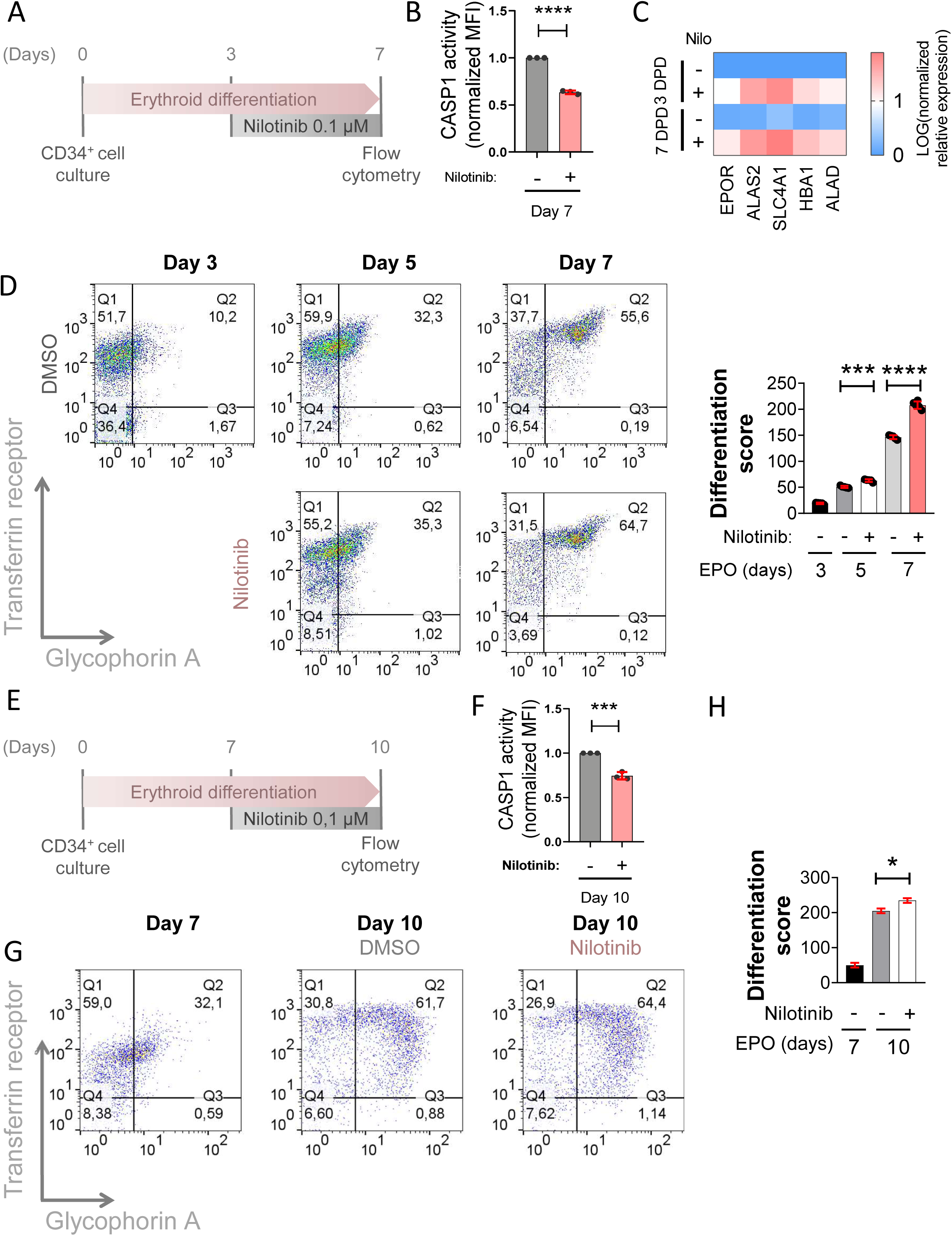
Nilotinib promotes erythropoiesis of human HSPCs from healthy donors. (A, E) Primary human CD34^+^/CD133^+^ HSPCs from healthy donors were cultured with EPO, and 0.1 µM nilotinib was added from days 3 to 7 (A) or days 7 to 11 (E). (B, F) CASP1 activity was measured by flow cytometry with FAM FLICA and normalized to untreated control cells. (D, G) Erythroid differentiation was assessed by flow cytometry after staining with anti-CD235A-APC (Glycophorin A) and anti-CD71-FITC (Transferrin Receptor). Representative dot plots of differentiation stages are shown. The differentiation score was calculated as the ratio between CD235A^+^/CD71^+^ (intermediate erythroid progenitors) and CD235A^-^/CD71^+^ (early erythroid progenitors). MFI, mean fluorescence intensity. Data are shown as the mean ± SEM.**p<0.01 and ****p<0.0001 according to a Student’s *t*-test.

### Several FDA/EMA-approved TKIs alleviate defective erythropoiesis in a zebrafish model of DBAS

Considering our previous results showing the impact of nilotinib over erythroid differentiation in human cells, we next tested whether other TKIs were able to show the same effects in zebrafish hematopoiesis. TKIs were developed for the treatment of Philadelphia chromosome (Ph)-positive chronic myeloid leukemia (CML), which is driven by the chimeric *BCR-ABL* gene. The first-generation drug imatinib showed an incredibly strong response in Ph-positive CML patients increasing their survival rate and life expectancy (Cortes et al., 2021; Kronick et al., 2023). Due to the resistance of imatinib during CML treatment, the second generation of TKIs were developed including bosutinib(Golas et al., 2003), dasatinib(Shah et al., 2004), nilotinib(Weisberg et al., 2005) and ponatinib(O’Hare et al., 2009). In our previous study, nilotinib robustly inhibited the Zaka/P38/Nlrp1 inflammasome in zebrafish larvae resulting in increased erythrocyte counts, decreased myeloid counts and reduced caspase-1 activity. We tested imatinib, dasatinib, ponatinib and bosutinib and none of the drugs affected development and survival (**Figures 3A, 3B**). Notably, all TKIs reduced caspase-1 activity and increased erythrocyte counts at the expense of neutrophils **(Figures 3C-3E**, **Figures 4A, 4C**). Strikingly, TKI treatment of the zebrafish Spint1a-deficient model, which is characterized by Nlrp1-driven neutrophilic inflammation(Rodriguez-Ruiz et al., 2023; Tyrkalska et al., 2019) (**Figure 4B**), alleviated neutrophilia without affecting neutrophil infiltration of inflamed skin (**Figure 4D**).

**Figure 3.**
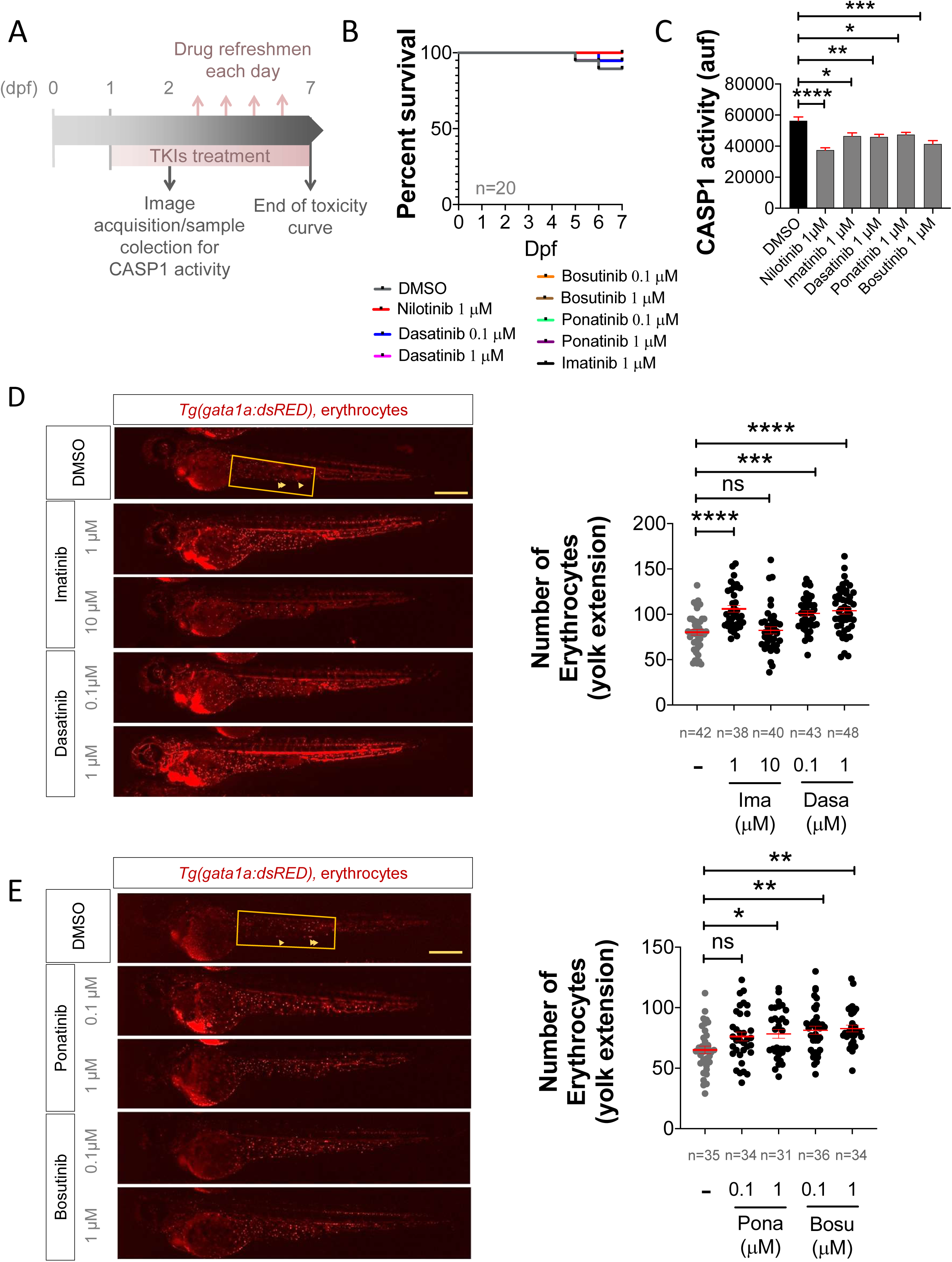
ZAKα inhibitors mimic the effects of nilotinib on hematopoiesis in zebrafish. (A) Schematic representation of the experimental design. Zebrafish larvae were treated with the indicated concentrations of different TKIs from 1 to 7 days post-fertilization (dpf). (B) Survival rates of zebrafish larvae after TKI treatment at the specified doses, monitored from 1 to 7 dpf.(C-E) Effects of TKIs on hematopoiesis in zebrafish larvae. Caspase-1 activity (C) and the number of erythrocytes (D, E) were evaluated in 2 dpf larvae treated via bath immersion with the indicated doses of TKIs from 1 to 2 dpf. Representative images of erythrocytes are shown. The region of interest is shown and erythrocytes labeled with arrowheads. Each dot represents one individual and the mean ± SEM for each group is also shown. P values were calculated using one-way ANOVA and Tukey’s multiple range test (C, D, E) or log rank test with Bonferroni correction (B). Data are shown as the means ± SEM of two technical replicates in (K). ns, non-significant; *P < 0.05, **P < 0.01, ***P<0.001 and ****P < 0.0001.

**Figure 4.**
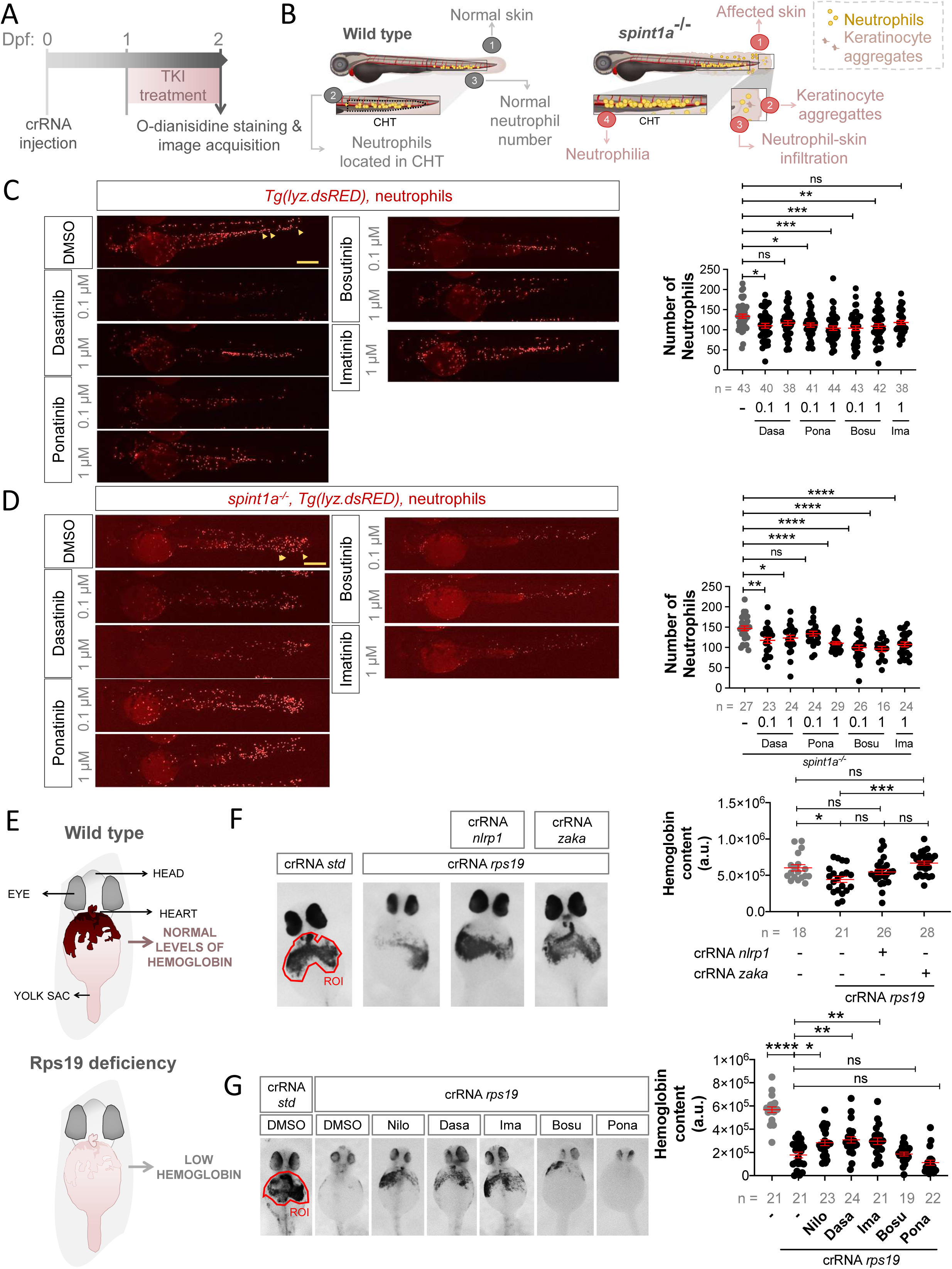
ZAKα Inhibitors Mimic the Effects of Nilotinib on Hematopoiesis in Zebrafish. (A, B, E) Schematic representations of the experimental designs. Zebrafish larvae were treated via bath immersion with the indicated TKIs from 1 to 2 days post-fertilization (dpf). (C, D) Quantification of neutrophils and representative images of treated larvae showing neutrophils (arrowheads). (F, G) Hemoglobin staining and quantification in Rps19-deficient zebrafish larvae generated by injecting embryos with gRNA/Cas9 complexes. Representative images of hemoglobin staining in treated larvae are shown. Regions of interest (ROI) are indicated in the images. Each dot represents one individual and the mean ± SEM for each group is also shown. P values were calculated using one-way ANOVA and Tukey’s multiple range test (C, D, F, G). ns, non-significant; *P < 0.05, **P < 0.01, ***P < 0.001 and ****p<0.0001.

These promising findings led us to explore the therapeutic potential of inhibiting the ZAKα/P38/NLRP1 inflammasome with TKIs in the context of DBAS. To this end, we generated a zebrafish model of DBAS through genetic disruption of *rps19* using CRISPR-Cas9 technology, as *RPS19* mutations are the most common genetic alterations in DBAS and are associated with defective erythropoiesis(Da Costa et al., 2020). Genetic experiments confirmed that the Zaka/Nlrp1 inflammasome drives anemia in *Rps19*-deficient larvae, as inhibition of either Nlrp1 or Zaka fully rescued the anemia in this model (**Figures 4A, 4E, 4F**). Remarkably, treatment with nilotinib, dasatinib, and imatinib, but not ponatinib and bosutinib, partially rescued the severe anemia observed in *Rps19*-deficient larvae, as assessed by the hemoglobin content (**Figures 4G**), highlighting the potential of these drugs as therapeutic options for DBAS.

### TKIs promote erythroid differentiation of K562 cells by inhibiting the ZAKα/P38/NLRP1 inflammasome and stabilizing GATA1 protein

Given that our initial validation in zebrafish larvae demonstrated that other TKIs played a significant role in hematopoiesis, with effects like those previously observed with nilotinib, we next investigated whether these drugs regulated hematopoiesis in human cells through the inhibition of the ZAKα/P38/NLRP1 inflammasome. Previously, we showed that erythroid differentiation of K562 cells with hemin resulted in the phosphorylation of ZAKα and P38, leading to NLRP1 inflammasome activation and GATA1 cleavage by CASP1 (Figure 1A). Notably, nilotinib blocked this activation and promoted GATA1 accumulation(Rodríguez-Ruiz et al., 2023). Consistent with these findings, we observed that the first-generation TKI imatinib, at 1 µM but not at 0.1 µM, induced GATA1 protein accumulation in K562 cells undergoing hemin-induced erythroid differentiation, while suppressing ZAKα/P38 phosphorylation. Similarly, both doses of dasatinib tested effectively replicated the effects of nilotinib (**Figures 5A, 5B**). Interestingly, although ponatinib and bosutinib treatments exhibited a dose-dependent inhibitory effect on ZAKα/P38 phosphorylation, all tested concentrations led to GATA1 protein accumulation and promoted erythroid differentiation, as evidenced by hemoglobin accumulation and the intense red color of the cell pellets (**Figures 5A, 5C**). Consistent with these results, inhibition of the ZAKα/P38 axis by the different TKIs impaired CASP1 activation during erythroid differentiation of K562 cells (**Figures 5A, 5D**). These findings suggest that GATA1 accumulation could be a consequence of CASP1 inhibition mediated by TKIs, since GATA1 is cleaved and inactivated by CASP1(Tyrkalska et al., 2019).

**Figure 5.**
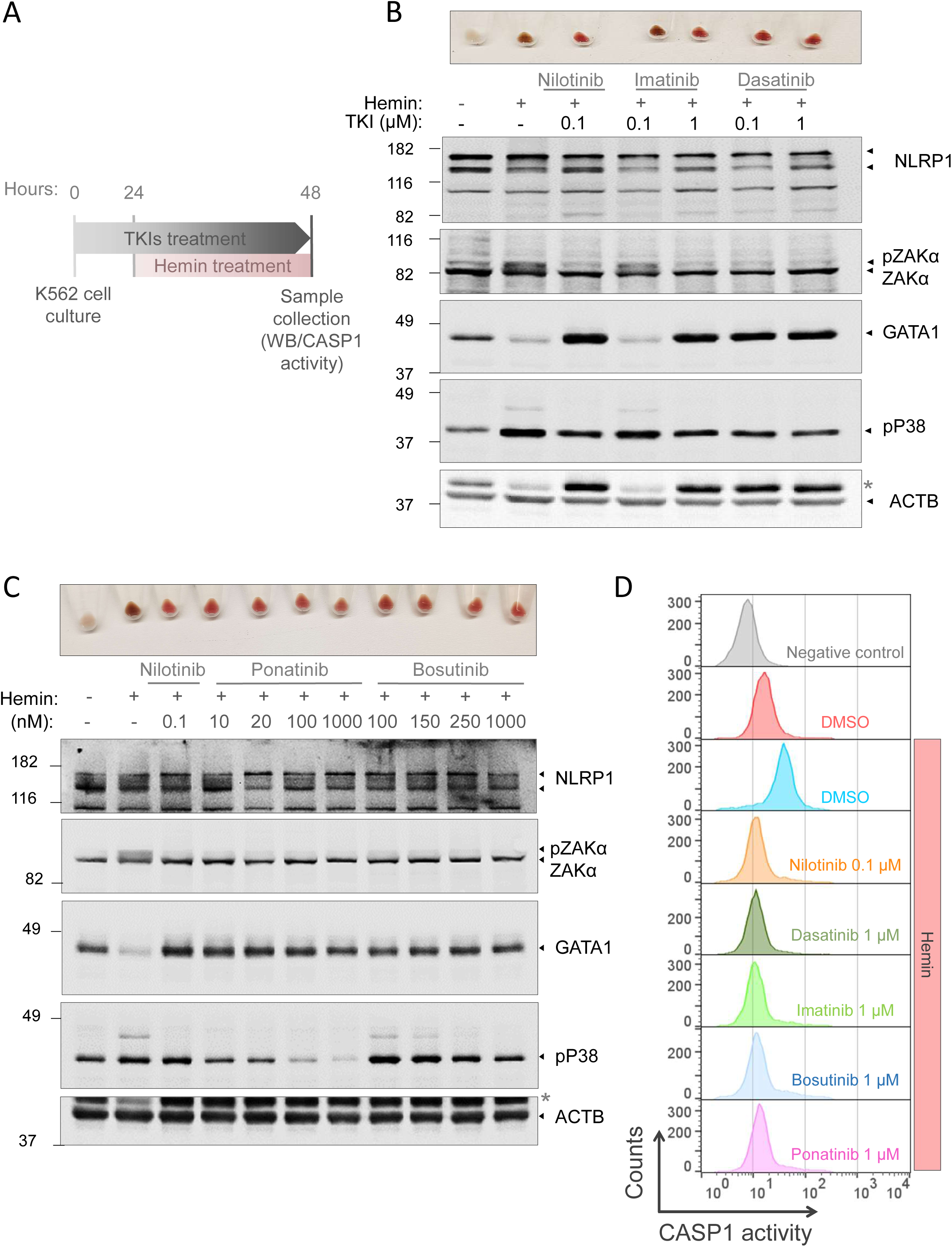
Inhibition of the ZAKα/P38/NLRP1 signaling pathway with TKIs facilitates terminal erythroid differentiation in K562 cells. (A) Schematic representation of the experimental design. (B-D) K562 cells were pretreated with 0.1 μM nilotinib (B-D), 0.1–1 μM imatinib, 0.1–1 μM dasatinib (B), 20–1000 nM ponatinib, or 100–1000 nM bosutinib (C) for 24 h and subsequently differentiated with 50 μM hemin for another 24 h. (B, C) Hemoglobin accumulation and protein levels of NLRP1, phosphorylated P38 (pP38), GATA1, ZAKα, and ACTB were assessed by Western blot. Immunoblots are representative of three independent experiments. The asterisk (*) denotes the GATA1 protein band. (D) CASP1 activity was quantified using FLICA-CASP1 staining and analyzed by flow cytometry. Representative histograms of CASP1 activity at different differentiation times are shown.

### Nilotinib alleviates defective erythroid differentiation of RPS19-deficient CD34^+^ HSPCs

To demonstrate the usefulness of nilotinib as a treatment for DBAS, we generated a DBAS model in human CD34^+^ cells by editing *RPS19* gene with CRISPR-Cas9 (Bhoopalan et al., 2023), reaching 35% of indel score after 13 days post-transfection (**Figure 6A**). RPS19 deficiency led to an increased accumulation of BFU-E (CD71⁺/CD235A⁻ cells) that failed to progress into erythroblasts (CD71⁺/CD235A⁺ cells), whereas nilotinib restored normal differentiation (**Figures 6B-D**). Strikingly, *RPS19* deficiency also led to enhanced CASP1 activity which was restored to normal levels by nilotinib (**Figure 6E**). Consistent with these findings, both imatinib (**Supplementary Figures 1A–1D, 2A-2D**) and dasatinib (**Supplementary Figures 1E–1K, 2E-2J**) replicated the effects of nilotinib on both wild-type and RPS19-deficient human CD34^+^ cells. Collectively, these results suggest that enhanced ribotoxic stress of RPS19-deficient erythroid progenitors results in hyperactivation of the ZAKα/P38/NLRP1/CASP1 inflammasome leading to exacerbated degradation of GATA1 and impaired erythroid differentiation, which can be effectively counteracted by TKIs such as nilotinib, imatinib and dasatinib.

**Figure 6.**
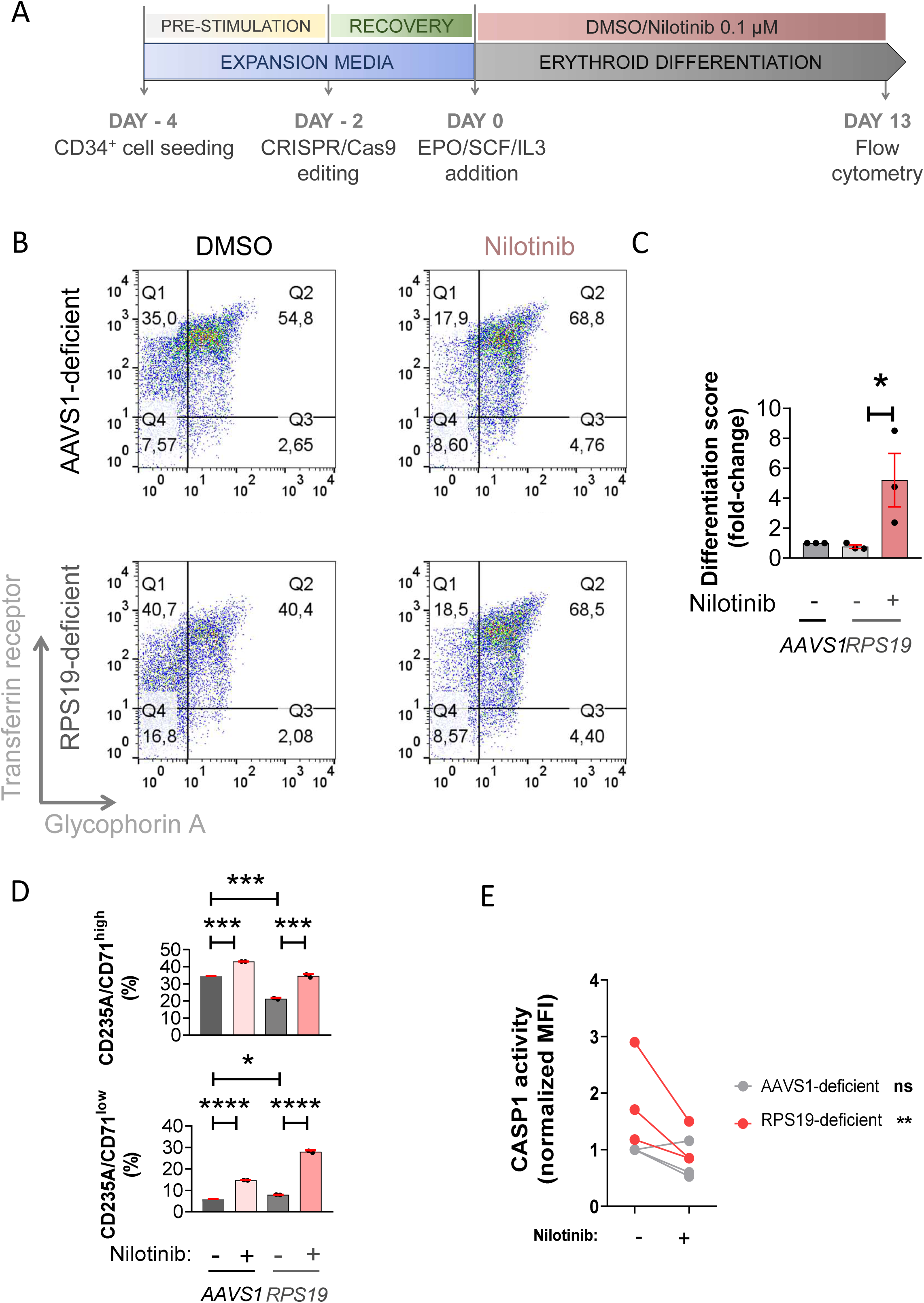
Nilotinib alleviates defective erythropoiesis of RPS19-deficient HSPCs. (A) Primary human CD34^+^ cells were purified from human cord blood or purchased from from ZenBio or StemCell Technologies, edited with CRISPR/Cas9 and differentiated for 13 days with EPO in the presence of either DMSO or 0.1 µM nilotinib. (B-D) Cells were stained with anti-CD235A-APC (Glycophorin A) and anti-CD71-FITC (Transferrin Receptor), or FAM FLICA, and erythroid differentiation (B-D) and CASP1 activity (E) were then analyzed by flow cytometry. Representative dot plots at different differentiation times are shown in B. The differentiation score was calculated as the ratio between CD235A^+^/CD71^+^ (intermediate erythroid progenitors) and CD235A^-^/CD71^+^ (early erythroid progenitors) (C), and the percentage of CD235A^+^/CD71^high^ (erythroblasts) and CD235A^+^/CD71^low^ (reticulocytes) (D) and CASP1 activity were determined at 13 days of culture (E). Data are shown as the mean ± SEM. P values were calculated using one-way ANOVA and Tukey’s multiple range test (C, D) or a Student’s *t*-test (E). ns, non-significant; *p<0.05; **p<0.01; ***p<0.01 and ****p<0.0001.

### TKIs alleviates defective erythroid differentiation of HSPCs from DBAS patients

We next examined whether the different TKIs tested in zebrafish, K562 cells and the *RSP19*-edited model of DBAS, were also able to alleviate defective erythropoiesis of HSPCs from DBAS patients. To do so, mononuclear cells from bone marrow aspirates (BMMCs) or peripheral blood (PBMCs) were isolated by isopycnic centrifugation and seeded in methylcellulose medium containing EPO and colony-stimulating factors (CSFs). The results showed that all TKI tested, but ponatinib, strongly increased the number of BFU-Es derived from HSPCs of DBAS patients (**Figure 7A-7G**). Importantly, although the effects of TKIs on the number of CFU-GMs varied greatly among patients and were not statistically significant overall, the highest dose of nilotinib tested (0.1 µM) was also able to significantly increase the number of myeloid colonies (**Figure 7H-7M**). These results suggest that the inhibition of the ZAKα/P38/NLRP1/CASP1 axis with TKIs may alleviate the impaired erythropoiesis of patients suffering from DBAS and, therefore, they are attractive to be repurposed for this disease.

**Figure 7.**
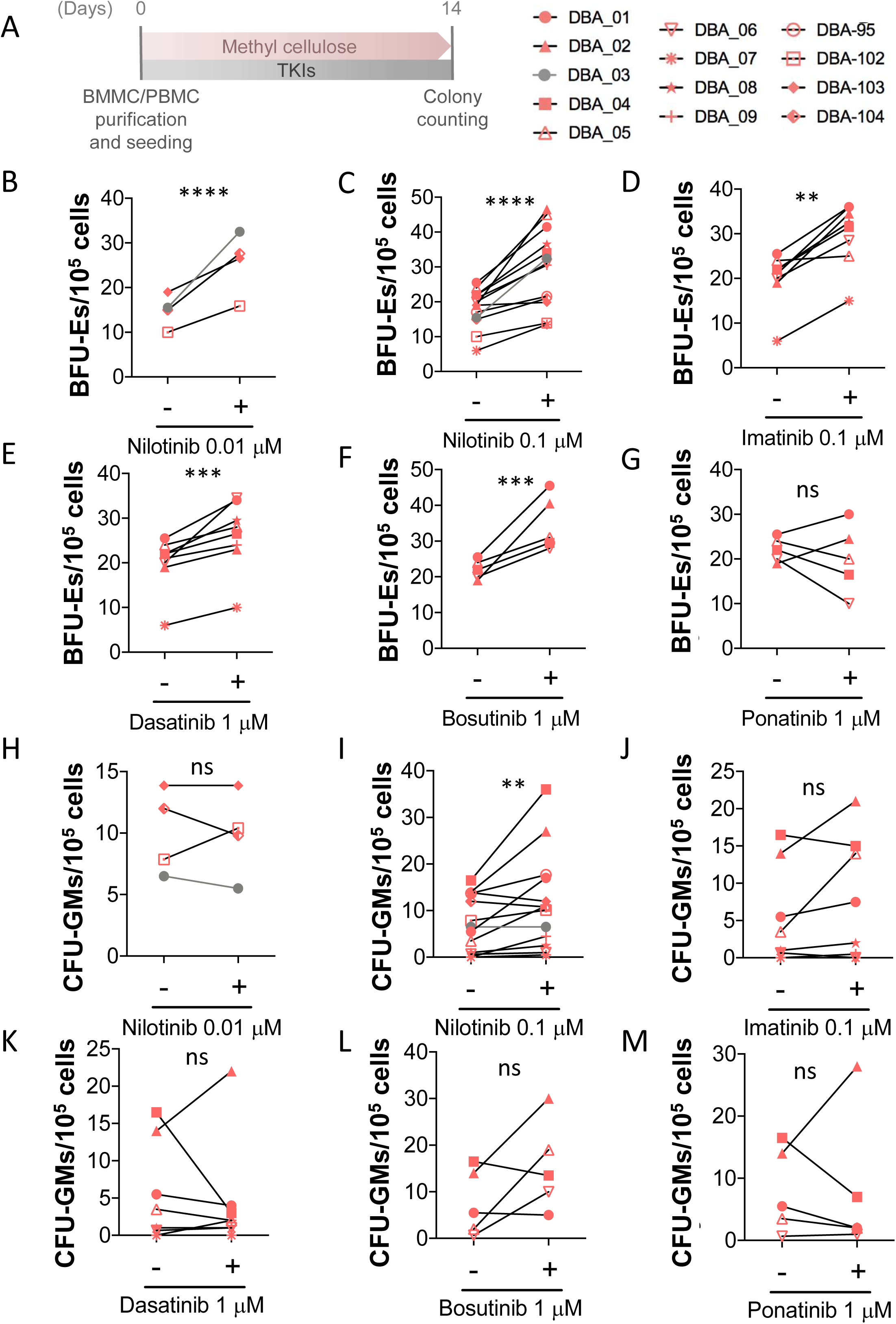
TKIs alleviate erythropoiesis defects of HSPCs from patients with DBAS. A) Peripheral blood mononuclear cells (PBMCs, red) and bone marrow mononuclear cells (BMMCs, grey) were isolated from 13 patients with DBAS. Cells were cultured for 2 weeks in human methylcellulose complete medium at 37 °C, with or without the indicated TKIs. Burst-forming unit-erythroid (BFU-E) and colony-forming unit-granulocyte macrophage (CFU-GM) colonies were counted based on standard morphological criteria. (B-M) Each dot represents data from an individual patient. The increase in erythroid colonies (BFU-E; panels B-G) and myeloid colonies (CFU-GM; panels H, M) upon treatment with TKIs is depicted. Data are shown as the mean ± SEM. ns, non-significant; **p<0.01, ***P<0.001 and ****P<0.0001 according to a Student’s *t*-test.

**Figure 8.**
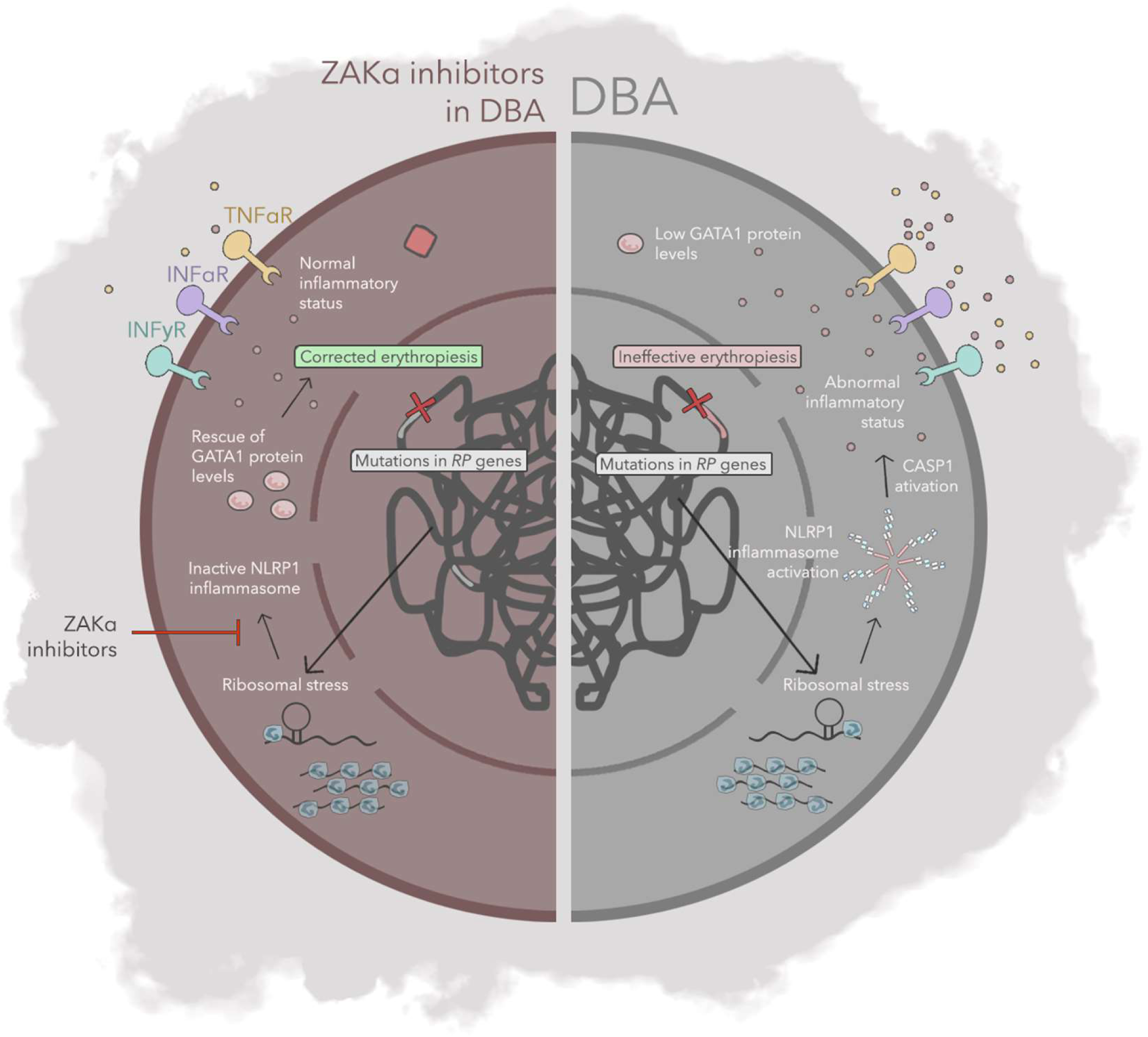
Working Model of ZAKα Inhibitors in DBA. Left: Mutations in ribosomal protein (RP) genes induce ribosomal stress, leading to impaired GATA1 translation and triggering assembly of the NLRP1 inflammasome. This, in turn, activates caspase-1, resulting in GATA1 degradation and defective erythropoiesis. Right: ZAKα inhibitors block the ribosomal stress-mediated activation of the NLRP1 inflammasome, thereby rescuing GATA1 protein levels and restoring normal erythropoiesis.

## Discussion

The importance of the inflammasome as therapeutic targets for the treatment of pathologies has gained interest in the clinic during the last years due to their recently reported strong implications in several diseases. Nowadays, most of FDA/EMA-approved drugs are designed to reach inflammasome-related targets, such as IL1B and IL18, instead of reaching directly the inflammasomes(Yao et al., 2024). Direct inhibition of specific inflammasomes components has also shown promising results in animal disease models. For instance, NLRP3 inhibitors, including MCC950, have been successfully used in Alzheimer’s disease by reducing memory deficits, inflammasome activity and the levels of Aβ plaques in APP/PS1 and TgCRND8 mice, meanwhile indirect inhibitors targeting either upstream or downstream proteins, such as the caspase-1 inhibitor VX-765 or the GSDMD inhibitor disulfiram decreased IL1B levels and improved cognitive abilities in 5×FAD mice (Ravichandran and Heneka, 2024). Tranilast, a drug that inhibits NLRP3 assembly by binding to its NATCH domain, approved to treat allergies, has shown a promising effect in vitiligo improving melanogenesis and melanosome translocation via attenuating IL1B secretion in keratinocytes (Zhuang et al., 2020). Caffeic acid phenethyl ester, which directly binds ASC, can be used to prevent NLRP3-ASC interactions triggered by monosodium urate (MSU) crystals and, therefore, has been proposed for the treatment of acute gout (Lee et al., 2016).

Recent studies, including some from our group, have extended the relevance of the inflammasome to hematological diseases (Ratajczak et al., 2020; Rodriguez-Ruiz et al., 2020). Our group initially reported that CASP1 was able to directly cleave and promote the degradation of the master erythroid transcription factor GATA1, finetuning directly GATA1 and indirectly SPI1 levels to control the myeloid-erythroid lineage decision of HSPCs(Tyrkalska et al., 2019). More recently, we identified the NLRP1 inflammasome being responsible for the activation of CASP1 in HSPCs, its strict negative regulation by LRRFIP1 and FLII in these cells, and its activation by phosphorylation of its linker domain by the ZAKα/P38 kinase axis upon ribosomal stress(Rodríguez-Ruiz et al., 2023). This discovery has revealed ZAKα as a druggable target using FDA/EMA approved TKIs for the treatment of congenital anemias, such as DBAS, where ribosome mutations lead to both ribosomal stress and inefficient GATA1 translation(Ludwig et al., 2014). Thus, we have shown that nilotinib was able to block the activation of NLRP1 inflammasome by ZAKα/P38 kinase signaling pathway resulting in reduced CASP1 activity, and enhanced GATA1 accumulation and erythroid differentiation of K562 cells. We extend here these observations by showing that treatment of K562 cells with nilotinib robustly promotes erythroid differentiation by increasing the transcript levels of genes encoding structural and functional genes required for erythropoiesis, such as those involved in iron acquisition, hemoglobin synthesis and erythrocyte structure. Given that these genes are regulated by GATA1, and that inhibition of the ZAKα/P38/NLRP1/CASP1 axis by nilotinib leads to increased GATA1 protein levels, it is likely that nilotinib induces their expression indirectly through GATA1 stabilization.

The usefulness of nilotinib, imatinib and dasatinib to be repurposed to treat DBAS was further confirmed using a *RPS19*-edited model of DBAS where they not only restored impaired erythroid differentiation of RPS19-deficient HSPCs but also exacerbated CASP1 activity, likely resulting from increased ribosomal stress(Khajuria et al., 2018). Furthermore, nilotinib, imatinib, dasatinib and bosutinib, all alleviated defective erythropoiesis of HSPCs from DBAS patients harboring different ribosome mutations, and phenocopied the effect of nilotinib in K562 cells and zebrafish; that is they (i) inhibited ZAKα/P38/NLRP1/CASP1 in K562 cells leading to increased GATA1 levels and erythroid differentiation, (ii) enhanced erythropoiesis at the expenses of granulopoiesis in zebrafish larvae, and (iii) ameliorated neutrophilia in a zebrafish model of neutrophilic inflammation. Therefore, these TKIs are promising drugs for the treatment of DBAS and likely other blood diseases characterized by defective erythropoiesis, such as Fanconi anemia, thalassemia, and myelodysplastic syndromes(Cappellini et al., 2023; Da Costa et al., 2020; Dorenkamp et al., 2023; Guerra et al., 2023) as well as myeloid lineage bias of chronic inflammatory diseases(Marzano et al., 2018; Weiss, 2015), cancer(Wu et al., 2014) and aging(Elias et al., 2017). Of note, although chronic myeloid leukemia (CML) patients treated with TKIs often exhibit anemia, this anemia is primarily attributable to the underlying disease rather than a direct effect of the drugs (Liu et al., 2020). Many CML patients present with anemia at diagnosis—approximately 35% in the chronic phase (Liu et al., 2020)—and early myelosuppression, affecting multiple blood lineages, is common following TKI initiation. This transient cytopenia reflects the suppression of the leukemic clone combined with pre-existing inhibition of normal hematopoiesis by malignant cells, with recovery typically observed once remission is achieved (Steegmann et al., 2016). Furthermore, imatinib is approved for gastrointestinal stromal tumors (GIST), a non-hematologic condition, where severe anemia (grade 3 or 4) occurs at very low rates (approximately 5.4% and 0.7%, respectively) and is likely related to bleeding complications rather than a direct drug effect (https://www.ema.europa.eu/en/documents/product-information/glivec-epar-product-information_en.pdf). These observations underscore that TKI-associated hematologic side effects are transient, dose-dependent, and reversible, further reinforcing their potential as therapeutic agents for congenital anemias.

In summary, the identification of the pivotal role of the NLRP1 inflammasome in hematopoiesis and its activation mechanisms has unveiled a promising therapeutic target for hematopoietic disorders, including congenital anemias and lineage bias disorders. Notably, the substantial efficacy of TKIs in enhancing erythroid differentiation in HSPCs from DBAS patients, as well as in the RPS19-edited model of DBAS, highlights these drugs as promising emergent treatments for this rare disease. Currently, available therapies for DBAS are limited to glucocorticoids, blood transfusions, hematopoietic stem cell transplantation, and gene therapy(Liu and Karlsson, 2023), underscoring the potential impact of TKIs as a novel therapeutic option.

## Acknowledgements

We thank I. Fuentes, P.J. Martínez, M. Ródenas and M.E. Rubio for their excellent technical assistance, and P. Crosier, H. and L.I. Zon for the zebrafish lines.

This work was supported by Fundación Séneca, Agencia de Ciencia y Tecnología de la Región de Murcia (research grants 21887/PI/22 and 22242/PDC/23 to VM), MCIN/AEI/10.13039/501100011033 (research grants 2020-113660RB-I00 to VM, and Juan de la Cierva-Incorporación postdoctoral contract to SDT), ISCIII (Miguel Servet CP21/00028 and CP23/00049 to DG-M and SDT, respectively), CIBERER (ER24P7AC7682-ACCI23-12-7682 to AM-L), Consejería de Salud-CARM (ZEBER contract to AM-L), and Diamond-Blackfan Anemia Foundation (USA). The funders had no role in the study design, data collection and analysis, decision to publish, or preparation of the manuscript.

## Author contributions

AM-L, SDT and VM conceived the study; JML-G, LR-R, MP, JP performed the research; JML-G, LR-R, MP, JP, SN, JLF, CB, AJ, LM-S, CDdH, GLdH, JZ, JS, MPS, MLC, DG-M, AM-L, SDT and VM analyzed data and provided materials and ideas; JML-G and VM wrote the original draft; VM edited the final version with minor contributions from all authors. All authors have read and agreed to the published version of the manuscript.

## Data availability

The RNA seq raw data will be made available at GEO upon acceptance. The rest of datasets generated and/or analyzed during the current study are available from the corresponding author (VM) on reasonable request.

---

**Supplementary Figure 1.**
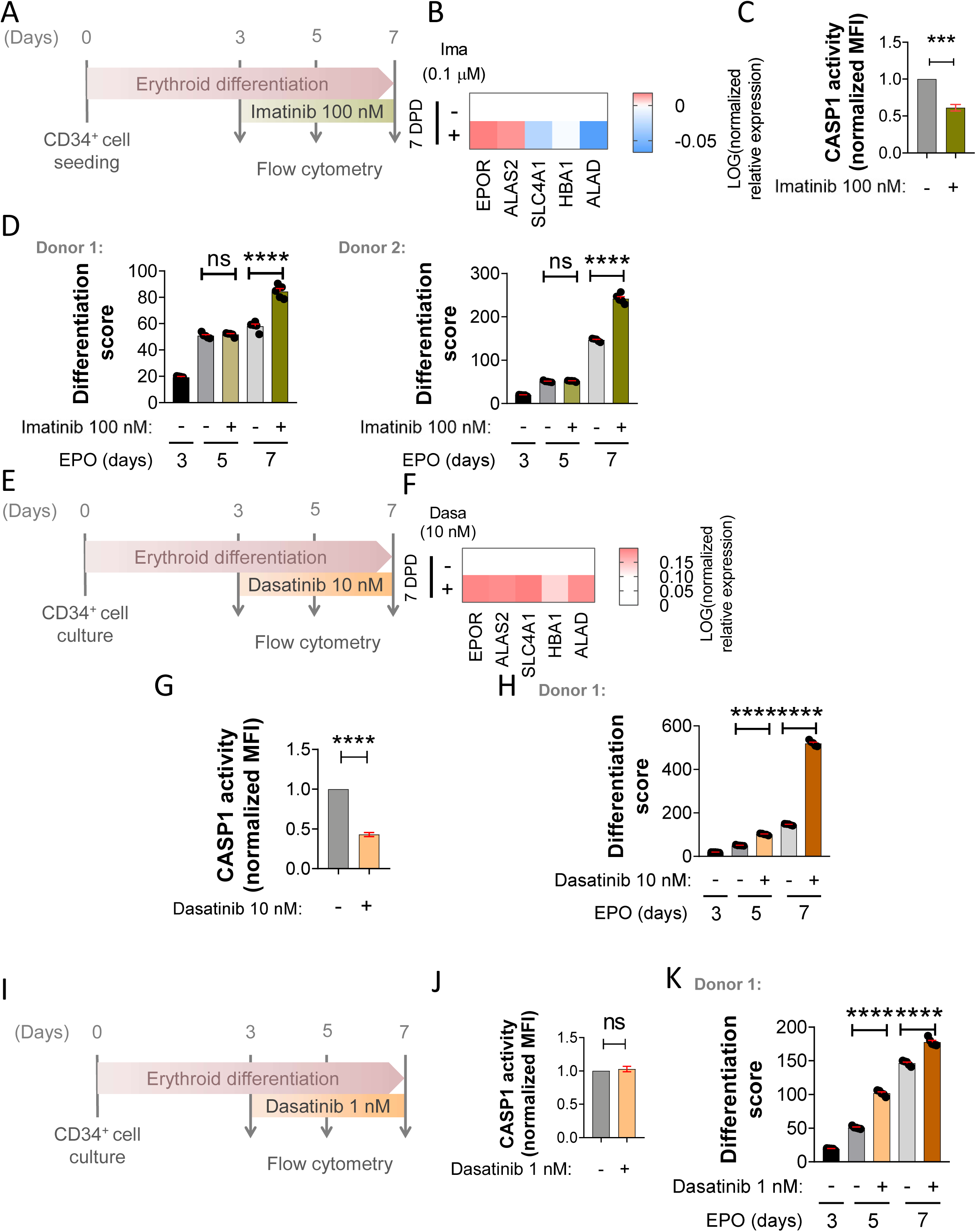
(related to Figs. 2 and 6). Imatinib and dasatinib replicate the effects of nilotinib on human HSPCs from healthy donors. Primary human CD34^+^ HSPCs from healthy donors were differentiated with EPO in the presence of 0.1 µM imatinib (A-D), or 10 (E-H) and 1 nM (I-K) dasatinib from 3 to 7 days of culture. Cells were stained with anti-CD235A-APC (Glycophorin A) and anti-CD71-FITC (Transferrin Receptor), and erythroid differentiation was then analyzed by flow cytometry. The transcript levels of GATA1-dependent genes (B), caspase-1 activity determined with FAM FLICA (D,G) and the differentiation score calculated as the ratio between CD235A^+^/CD71^+^ (intermediate erythroid progenitors) and CD235A^-^/CD71^+^ (early erythroid progenitors) (D, H, K) at 7 dpd are shown. Data are shown as the mean ± SEM. P values were calculated using one-way ANOVA and Tukey’s multiple range test (D, H, K) or a Student’s *t*-test (C, G). ns, non-significant; *p<0.05; **p<0.01; ***p<0.01 and ****p<0.0001.

**Supplementary Figure 2.**
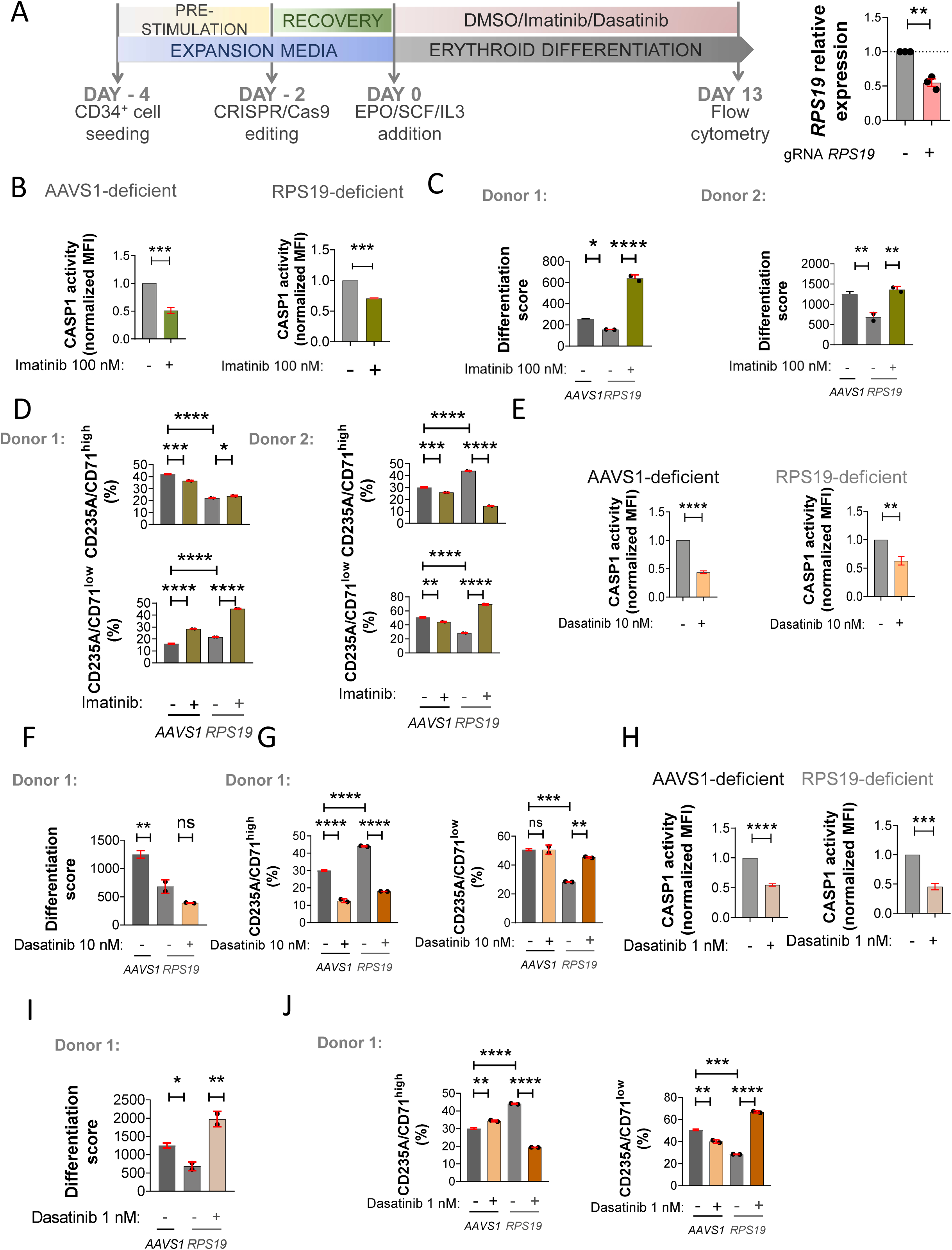
(related to Figs. 2 and 6). Imatinib and dasatinib alleviate defective erythropoiesis of RPS19-deficient HSPCs. (A) Primary human CD34^+^ cells were purchased from from ZenBio or StemCell Technologies, edited with CRISPR/Cas9 and differentiated for 13 days with EPO in the presence of of 0.1 µM imatinib (A-D) or 10 nM dasatinib (F-J) from 3 to 13 days of culture. (B-D). The transcript levels of RPS19 were analyzed by RT-qPCR at 13 days of culture (A). Cells were stained with either FAM FLICA or anti-CD235A-APC (Glycophorin A) and anti-CD71-FITC (Transferrin Receptor), and CASP1 activity (B, H), erythroid differentiation (B-D, F, G, I, J) were then analyzed by flow cytometry. The differentiation score was calculated as the ratio between CD235A^+^/CD71^+^ (intermediate erythroid progenitors) and CD235A^-^/CD71^+^ (early erythroid progenitors) (C, F, I), and the percentage of CD235A^+^/CD71^high^ (erythroblasts) and CD235A^+^/CD71^low^ (reticulocytes) (D, G, J) at 13 days of culture (E). Data are shown as the mean ± SEM. P values were calculated using one-way ANOVA and Tukey’s multiple range test (C, D, F, G, I, J) or a Student’s *t*-test (A, E, H). ns, non-significant; *p<0.05; **p<0.01; ***p<0.01 and ****p<0.0001.

**Supplementary Table 1.**
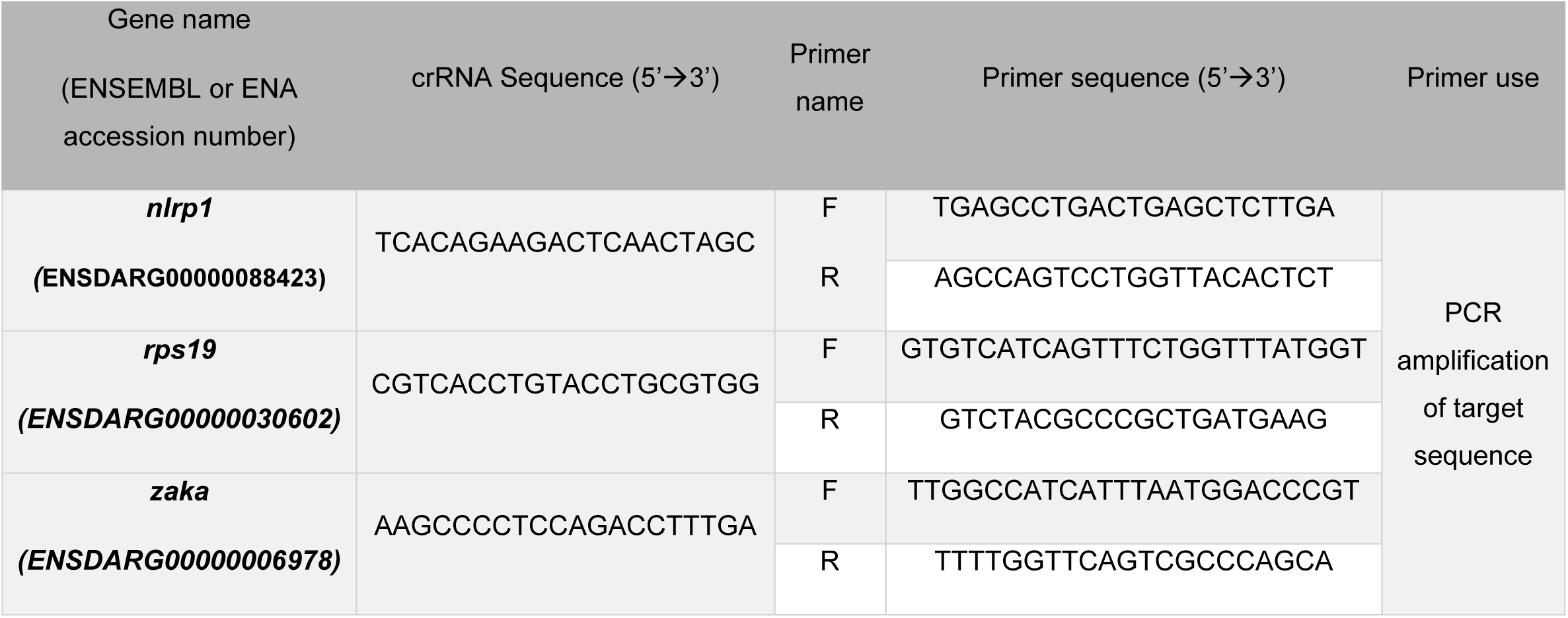
Genetic tools used for CRISPR/Cas9 experiments with zebrafish embryo. The gene symbols followed the Zebrafish Nomenclature Guidelines (http://zfin.org/zf_info/nomen.html).

**Supplementary Table 2.** List of mutations of each DBA patient who participated in this study.

| **Patient ID** | **Genetic variant** | **Gender** |
| --- | --- | --- |
| **DBA_01** | *RPS10: c.268_275del,*<br>*p.Val90HisfsTer5* | *Female* |
| **DBA_02** | *RPS26: c.328+1G>T* | *Male* |
| **DBA_03** | *RPS19: c.356+1G>T* | *Female* |
| **DBA_04** | *RPS10: c.337C>T, p.Arg113Ter* | *Female* |
| **DBA_05** | *RPL11: c.6+2T>C* | *Male* |
| **DBA_06** | *RPL11: c.372_385del,*<br>*p.Ile125LeufsTer7* | *Female* |
| **DBA_07** | *RPL35A: c.82_84insCTT* | *Male* |
| **DBA_08** | *RPS19: c.388_389del,*<br>*p.Asp130SerfsTer23* | *Male* |
| **DBA_09** | *RPL5: c.175_176delGA,*<br>*p.Asp59TyrfsTer53* | *Female* |
| **DBA_95** | *RPS19: c.172+1G>A* | *No data* |
| **DBA_102** | *RPS19: Mutation unavailable* | *No data* |
| **DBA_103** | *RPS26: c.131_132del,*<br>*p.Ile44SerfsTer11* | *No data* |
| **DBA_104** | *RPL11: Mutation unavailable* | *No data* |

**Supplementary Table 3.**
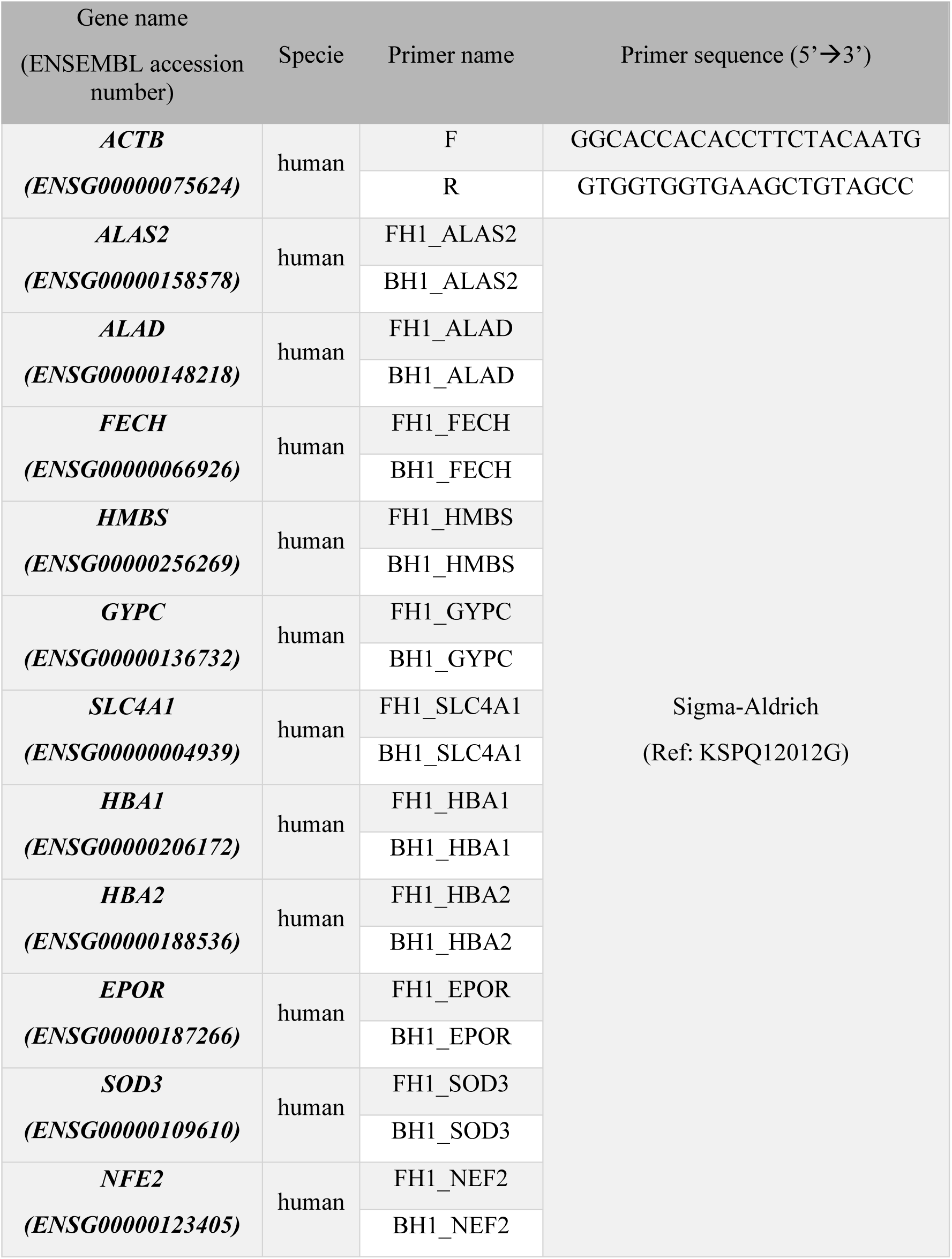
Primers used for RT-qPCR.

## References

Adamiak, M., A. Abdel-Latif, K. Bujko, A. Thapa, K. Anusz, M. Tracz, K. Brzezniakiewicz-Janus, J. Ratajczak, M. Kucia, and M.Z. Ratajczak. 2020. Nlrp3 Inflammasome Signaling Regulates the Homing and Engraftment of Hematopoietic Stem Cells (HSPCs) by Enhancing Incorporation of CXCR4 Receptor into Membrane Lipid Rafts. Stem Cell Rev Rep 16:954–967.

Angosto, D., G. López-Castejón, A. López-Muñoz, M.P. Sepulcre, M. Arizcun, J. Meseguer, and V. Mulero. 2012. Evolution of inflammasome functions in vertebrates: Inflammasome and caspase-1 trigger fish macrophage cell death but are dispensable for the processing of IL-1β. Innate Immun 18:815–824.

Bhoopalan, S.V., J.S. Yen, T. Mayuranathan, K.D. Mayberry, Y. Yao, M.A. Lillo Osuna, Y. Jang, J.S. Liyanage, L. Blanc, S.R. Ellis, M.W. Wlodarski, and M.J. Weiss. 2023. An RPS19-edited model for Diamond-Blackfan anemia reveals TP53-dependent impairment of hematopoietic stem cell activity. JCI Insight 8:

Cappellini, M.D., A.T. Taher, A. Verma, F. Shah, and O. Hermine. 2023. Erythropoiesis in lower-risk myelodysplastic syndromes and beta-thalassemia. Blood Rev 59:101039.

Cortes, J., C. Pavlovsky, and S. Saußele. 2021. Chronic myeloid leukaemia. The Lancet 398:1914–1926.

Da Costa, L., T. Leblanc, and N. Mohandas. 2020. Diamond-Blackfan anemia. Blood 136:1262–1273.

Dorenkamp, M., N. Porret, M. Diepold, and A. Rovó. 2023. A De Novo Frameshift Mutation in RPL5 with Classical Phenotype Abnormalities and Worsening Anemia Diagnosed in a Young Adult-A Case Report and Review of the Literature. Medicina (Kaunas) 59:

Elias, H.K., D. Bryder, and C.Y. Park. 2017. Molecular mechanisms underlying lineage bias in aging hematopoiesis. Semin Hematol 54:4–11.

Evavold, C.L., and J.C. Kagan. 2019. Inflammasomes: Threat-Assessment Organelles of the Innate Immune System. Immunity 51:609–624.

Frame, J.M., T. Long, C. Schuster-Kubaczka, V. Esain, S.E. Lim, G.Q. Daley, and T. North. 2018. Inflammasome-Mediated Regulation of Hematopoiesis in the Vertebrate Embryo. Blood 132:330–330.

Golas, J.M., K. Arndt, C. Etienne, J. Lucas, D. Nardin, J. Gibbons, P. Frost, F. Ye, D.H. Boschelli, and F. Boschelli. 2003. SKI-606, a 4-anilino-3-quinolinecarbonitrile dual inhibitor of Src and Abl kinases, is a potent antiproliferative agent against chronic myelogenous leukemia cells in culture and causes regression of K562 xenografts in nude mice. Cancer research 63:375–381.

Guerra, A., H. Parhiz, and S. Rivella. 2023. Novel potential therapeutics to modify iron metabolism and red cell synthesis in diseases associated with defective erythropoiesis. Haematologica 108:2582–2593.

Hall, C., M.V. Flores, T. Storm, K. Crosier, and P. Crosier. 2007. The zebrafish lysozyme C promoter drives myeloid-specific expression in transgenic fish. BMC Dev Biol 7:42.

He, Y., H. Hara, and G. Núñez. 2016. Mechanism and Regulation of NLRP3 Inflammasome Activation. Trends in Biochemical Sciences 41:1012–1021.

Khajuria, R.K., M. Munschauer, J.C. Ulirsch, C. Fiorini, L.S. Ludwig, S.K. McFarland, N.J. Abdulhay, H. Specht, H. Keshishian, D.R. Mani, M. Jovanovic, S.R. Ellis, C.P. Fulco, J.M. Engreitz, S. Schutz, J. Lian, K.W. Gripp, O.K. Weinberg, G.S. Pinkus, L. Gehrke, A. Regev, E.S. Lander, H.T. Gazda, W.Y. Lee, V.G. Panse, S.A. Carr, and V.G. Sankaran. 2018. Ribosome Levels Selectively Regulate Translation and Lineage Commitment in Human Hematopoiesis. Cell 173:90–103 e119.

Kronick, O., X. Chen, N. Mehra, A. Varmeziar, R. Fisher, D. Kartchner, V. Kota, and C.S. Mitchell. 2023. Hematological Adverse Events with Tyrosine Kinase Inhibitors for Chronic Myeloid Leukemia: A Systematic Review with Meta-Analysis. Cancers (Basel) 15:

Lee, H.E., G. Yang, N.D. Kim, S. Jeong, Y. Jung, J.Y. Choi, H.H. Park, and J.Y. Lee. 2016. Targeting ASC in NLRP3 inflammasome by caffeic acid phenethyl ester: a novel strategy to treat acute gout. Scientific Reports 6:38622.

Lenkiewicz, A.M., M. Adamiak, A. Thapa, K. Bujko, D. Pedziwiatr, A.K. Abdel-Latif, M. Kucia, J. Ratajczak, and M.Z. Ratajczak. 2019. The Nlrp3 Inflammasome Orchestrates Mobilization of Bone Marrow-Residing Stem Cells into Peripheral Blood. Stem Cell Rev Rep 15:391–403.

Li, H., F. Hu, R.P. Gale, M.A. Sekeres, and Y. Liang. 2022. Myelodysplastic syndromes. Nature Reviews Disease Primers 8:74.

Liu, Y., and S. Karlsson. 2023. Perspectives of current understanding and therapeutics of Diamond-Blackfan anemia. Leukemia

Liu, Z., Y. Shi, Z. Yan, Z. He, B. Ding, S. Tao, Y. Li, L. Yu, and C. Wang. 2020. Impact of anemia on the outcomes of chronic phase chronic myeloid leukemia in TKI era. Hematology 25:181–185.

Lopez-Castejon, G., M.P. Sepulcre, I. Mulero, P. Pelegrin, J. Meseguer, and V. Mulero. 2008. Molecular and functional characterization of gilthead seabream Sparus aurata caspase-1: the first identification of an inflammatory caspase in fish. Mol Immunol 45:49–57.

Ludwig, L.S., H.T. Gazda, J.C. Eng, S.W. Eichhorn, P. Thiru, R. Ghazvinian, T.I. George, J.R. Gotlib, A.H. Beggs, C.A. Sieff, H.F. Lodish, E.S. Lander, and V.G. Sankaran. 2014. Altered translation of GATA1 in Diamond-Blackfan anemia. Nature Medicine 20:748–753.

Martin, B.N., C. Wang, C.J. Zhang, Z. Kang, M.F. Gulen, J.A. Zepp, J. Zhao, G. Bian, J.S. Do, B. Min, P.G. Pavicic, Jr., C. El-Sanadi, P.L. Fox, A. Akitsu, Y. Iwakura, A. Sarkar, M.D. Wewers, W.J. Kaiser, E.S. Mocarski, M.E. Rothenberg, A.G. Hise, G.R. Dubyak, R.M. Ransohoff, and X. Li. 2016. T cell-intrinsic ASC critically promotes T(H)17-mediated experimental autoimmune encephalomyelitis. Nat Immunol 17:583–592.

Martinon, F., K. Burns, and J. Tschopp. 2002. The inflammasome: a molecular platform triggering activation of inflammatory caspases and processing of proIL-beta. Mol Cell 10:417–426.

Marzano, A.V., A. Borghi, D. Wallach, and M. Cugno. 2018. A Comprehensive Review of Neutrophilic Diseases. Clin Rev Allergy Immunol 54:114–130.

O’Hare, T., W.C. Shakespeare, X. Zhu, C.A. Eide, V.M. Rivera, F. Wang, L.T. Adrian, T. Zhou, W.S. Huang, Q. Xu, C.A. Metcalf, 3rd, J.W. Tyner, M.M. Loriaux, A.S. Corbin, S. Wardwell, Y. Ning, J.A. Keats, Y. Wang, R. Sundaramoorthi, M. Thomas, D. Zhou, J. Snodgrass, L. Commodore, T.K. Sawyer, D.C. Dalgarno, M.W. Deininger, B.J. Druker, and T. Clackson. 2009. AP24534, a pan-BCR-ABL inhibitor for chronic myeloid leukemia, potently inhibits the T315I mutant and overcomes mutation-based resistance. Cancer Cell 16:401–412.

Pfaffl, M.W. 2001. A new mathematical model for relative quantification in real-time RT-PCR. Nucleic Acids Res 29:e45.

Phan, T.G., I. Grigorova, T. Okada, and J.G. Cyster. 2007. Subcapsular encounter and complement-dependent transport of immune complexes by lymph node B cells. Nature Immunology 8:992–1000.

Ratajczak, M.Z., M. Adamiak, A. Thapa, K. Bujko, K. Brzezniakiewicz-Janus, and A.M. Lenkiewicz. 2019. NLRP3 inflammasome couples purinergic signaling with activation of the complement cascade for the optimal release of cells from bone marrow. Leukemia 33:815–825.

Ratajczak, M.Z., K. Bujko, M. Cymer, A. Thapa, M. Adamiak, J. Ratajczak, A.K. Abdel-Latif, and M. Kucia. 2020. The Nlrp3 inflammasome as a “rising star” in studies of normal and malignant hematopoiesis. Leukemia 34:1512–1523.

Ravichandran, K.A., and M.T. Heneka. 2024. Inflammasomes in neurological disorders - mechanisms and therapeutic potential. Nat Rev Neurol

Rodriguez-Ruiz, L., J.M. Lozano-Gil, C. Lachaud, P. Mesa-Del-Castillo, M.L. Cayuela, D. Garcia-Moreno, A.B. Perez-Oliva, and V. Mulero. 2020. Zebrafish Models to Study Inflammasome-Mediated Regulation of Hematopoiesis. Trends Immunol 41:1116–1127.

Rodriguez-Ruiz, L., J.M. Lozano-Gil, E. Naranjo-Sanchez, E. Martinez-Balsalobre, A. Martinez-Lopez, C. Lachaud, M. Blanquer, T.K. Phung, D. Garcia-Moreno, M.L. Cayuela, S.D. Tyrkalska, A.B. Perez-Oliva, and V. Mulero. 2023. ZAKalpha/P38 kinase signaling pathway regulates hematopoiesis by activating the NLRP1 inflammasome. EMBO Mol Med 15:e18142.

Rodríguez-Ruiz, L., J.M. Lozano-Gil, E. Naranjo-Sánchez, E. Martínez-Balsalobre, A. Martínez-López, C. Lachaud, M. Blanquer, T.K. Phung, D. García-Moreno, M.L. Cayuela, S.D. Tyrkalska, A.B. Pérez-Oliva, and V. Mulero. 2023. ZAKα/P38 kinase signaling pathway regulates hematopoiesis by activating the NLRP1 inflammasome. EMBO Mol Med 15:e18142.

Shah, N.P., C. Tran, F.Y. Lee, P. Chen, D. Norris, and C.L. Sawyers. 2004. Overriding imatinib resistance with a novel ABL kinase inhibitor. Science 305:399–401.

Smith, R.D., J.D. Malley, and A.N. Schechter. 2000. Quantitative analysis of globin gene induction in single human erythroleukemic cells. Nucleic Acids Res 28:4998–5004.

Steegmann, J.L., M. Baccarani, M. Breccia, L.F. Casado, V. Garcia-Gutierrez, A. Hochhaus, D.W. Kim, T.D. Kim, H.J. Khoury, P. Le Coutre, J. Mayer, D. Milojkovic, K. Porkka, D. Rea, G. Rosti, S. Saussele, R. Hehlmann, and R.E. Clark. 2016. European LeukemiaNet recommendations for the management and avoidance of adverse events of treatment in chronic myeloid leukaemia. Leukemia 30:1648–1671.

Traver, D., B.H. Paw, K.D. Poss, W.T. Penberthy, S. Lin, and L.I. Zon. 2003. Transplantation and in vivo imaging of multilineage engraftment in zebrafish bloodless mutants. Nat Immunol 4:1238–1246.

Tyrkalska, S.D., S. Candel, D. Angosto, V. Gómez-Abellán, F. Martín-Sánchez, D. García-Moreno, R. Zapata-Pérez, Á. Sánchez-Ferrer, M.P. Sepulcre, P. Pelegrín, and V. Mulero. 2016. Neutrophils mediate Salmonella Typhimurium clearance through the GBP4 inflammasome-dependent production of prostaglandins. Nat Commun 7:12077.

Tyrkalska, S.D., A.B. Perez-Oliva, L. Rodriguez-Ruiz, F.J. Martinez-Morcillo, F. Alcaraz-Perez, F.J. Martinez-Navarro, C. Lachaud, N. Ahmed, T. Schroeder, I. Pardo-Sanchez, S. Candel, A. Lopez-Munoz, A. Choudhuri, M.P. Rossmann, L.I. Zon, M.L. Cayuela, D. Garcia-Moreno, and V. Mulero. 2019. Inflammasome Regulates Hematopoiesis through Cleavage of the Master Erythroid Transcription Factor GATA1. Immunity 51:50–63 e55.

Weisberg, E., P.W. Manley, W. Breitenstein, J. Brüggen, S.W. Cowan-Jacob, A. Ray, B. Huntly, D. Fabbro, G. Fendrich, and E. Hall-Meyers. 2005. Characterization of AMN107, a selective inhibitor of native and mutant Bcr-Abl. Cancer cell 7:129–141.

Weiss, G. 2015. Anemia of Chronic Disorders: New Diagnostic Tools and New Treatment Strategies. Semin Hematol 52:313–320.

Westerfield, M. 2000. The Zebrafish Book. A Guide for the Laboratory Use of Zebrafish Danio* (Brachydanio) rerio. University of Oregon Press., Eugene, OR.

White, R.M., A. Sessa, C. Burke, T. Bowman, J. LeBlanc, C. Ceol, C. Bourque, M. Dovey, W. Goessling, C.E. Burns, and L.I. Zon. 2008. Transparent adult zebrafish as a tool for in vivo transplantation analysis. Cell Stem Cell 2:183–189.

Wu, W.C., H.W. Sun, H.T. Chen, J. Liang, X.J. Yu, C. Wu, Z. Wang, and L. Zheng. 2014. Circulating hematopoietic stem and progenitor cells are myeloid-biased in cancer patients. Proc Natl Acad Sci U S A 111:4221–4226.

Yao, J., K. Sterling, Z. Wang, Y. Zhang, and W. Song. 2024. The role of inflammasomes in human diseases and their potential as therapeutic targets. Signal Transduct Target Ther 9:10.

Zhuang, T., S. Li, X. Yi, S. Guo, Y. Wang, J. Chen, L. Liu, Z. Jian, T. Gao, P. Kang, and C. Li. 2020. Tranilast Directly Targets NLRP3 to Protect Melanocytes From Keratinocyte-Derived IL-1β Under Oxidative Stress. Front Cell Dev Biol 8:588.

